# iTissue

**DOI:** 10.1101/2025.06.04.657816

**Authors:** Carme Zambrana, Noël Malod-Dognin, Nataša Pržulj

## Abstract

Tumors are highly heterogeneous tissues, and this heterogeneity impacts metastasis and its progression. Spatial data are essential for studying these hetrogeneity. However, its limited resolution makes crucial the integration of spatial data with molecular networks to identify relevant genes for the progression of metastasis. We develop a new data integration method, named iTissue, based on Graph Non-negative Matrix Tri-Factorization that integrates spatial and single-cell transcriptomics with molecular interaction networks, producing low-dimensional space. For three clear cell renal cell carcinoma (ccRCC) patients, we applied iTissue to their tumour-core and tumour-normal interface samples. We identified the top 100 with the highest euclidean distance between these two samples, since we show that the genes with the highest euclidean distance are relevant for ccRCC. By taking the intersection of the top 100 genes for the three patients, we identify 50 genes enriched in GO-BP terms relevant for metastasis. Moreover, we find that we can cluster the three regions of the tumour-normal interface (healthy tissue, tumour tissue and interface) using the identified genes. Our framework is general and can enable insight into other diseases.

## Introduction

The tumor microenvironment (TME) plays a crucial role in the progression of metastasis. Within this microenvironment, factors such as extracellular matrix components, stromal cells, immune cells, and signaling molecules interact in complex ways to promote tumor growth, invasion, and the spread of cancer to distant sites. Spatial data are essential for studying the TME in the context of metastasis, as they allow a deeper understanding of the complex interactions between tumor cells and their surroundings. Unlike traditional bulk data, which provide an averaged picture of molecular changes, spatially resolved data allows to map the precise localization of key components within the TME. However, spatial data resolution is limited; thus the integration of these data with molecular networks is crucial to identify relevant genes for the progression of metastasis.

Matrix factorization methods, such as NMF and its extension Non-Negative Matrix Tri-Factorization (NMTF) and Graph-regulirased NMTF (GNMTF), are highly effective for integrating expression data with biological networks because they provide a robust framework to combine heterogeneous data types. Recently, NMTF has successfully been used to integrate single-cell transcriptomics data with molecular networks in the context of Parkinson Disease^1,2^. One of the strengths of NMTF-based methods is its ability to integrate high-dimensional data from different sources (e.g., gene expression, protein interactions, spatial localization) and project them into lower-dimensional embedding (latent) spaces^3^. In contrast to other artificial intelligence algorithms, these methods are linear in nature and define latent spaces with natural biological interpretations^4,5^. Moreover, the GNMTF extension allows to add information about any data type using graph regularisation^6^, which can be used to add the spatial dependencies of the spatial data. Thus, we propose iTissue, a data integration framework based on GNMTF to integrate spatial and single-cell transcriptomics data with molecular networks to uncover relevant genes for the progression of metastasis.

We use iTissue to study the difference between the tumour-core and tumour-normal interface of clear cell renal cell carcinoma (ccRCC) patients. First, for each patient and sample (tumour-core and tumour-normal interface), we integrate spatial and single-cell transcriptomics data with molecular networks and obtain the genes embeddings in the low-dimensional space created by the integration. Then, for each patient, we obtained the top 100 genes with the highest euclidean distance between the gene embeddings for the tumour-core and tumour-normal interface samples, since we show that the genes with the highest euclidean distance are relevant for ccRCC. From these top 100 genes, we identify 50 genes that are shared between the 3 patients and validate that they are relevant for metastatic mechanisms. Finally, we find that we can cluster the three regions of the tumour-normal interface (healthy tissue, tumour tissue and interface) using the identified genes.

## Results

We create our data fusion framework iTissue, based on graph-regularized non-negative matrix tri-factorization (GNMTF) to integrate spatial and single-cell transcriptomics data with molecular networks (i.e., protein-protein interactions (PPI), genetic interactions (GI), co-expression (CoEx) and metabolic interactions (MI)). To add the spatial dependencies of the spatial data, we use graph regularisation penalties to favour embedding close in space the spots that are neighbours in the spatial spot graph constructed by connecting each spot to all its neighbours (see figure 8; for more details on the data that we used see “Data collection” section and on our framework see “iTissue data integration framework” section).

### Ablation study

To assess the contribution of each molecular network (PPI, GI, CoEx and MI) to create a gene space that is functionally organised, we integrate the spatial and single-cell data with the different possible combinations of these four networks using our iTissue data integration framework (for more details, see section “iTissue data integration framework”). For each network, we used two versions: its adjacency matrix and a weighted version of it (for more details, see section “Molecular networks”). To measure the functional organisation of the space, we cluster the genes and compute the enrichment of these clusters in the following biological annotations: Gene Ontology-Biological Processes (GO-BP)^7^, Reactome^8^ and KEGG^9^ (for more details, see section “Extracting clusters of genes” and “Enrichment Analysis”). For the three biological annotations, the weighted version of all combinations of networks, except GI+CoEx and GI+MI, obtained higher percentage of enriched genes than the adjacency matrix counterparts (see Figure 2). The combination of PPI and MI networks obtained the highest percentage of enriched genes for Reactome and KEGG pathways, and the second-best percentage for GO-BP. Thus, we will use the combination of spatial and single-cell transcriptomic and PPI and MI networks.

**Figure 1.**
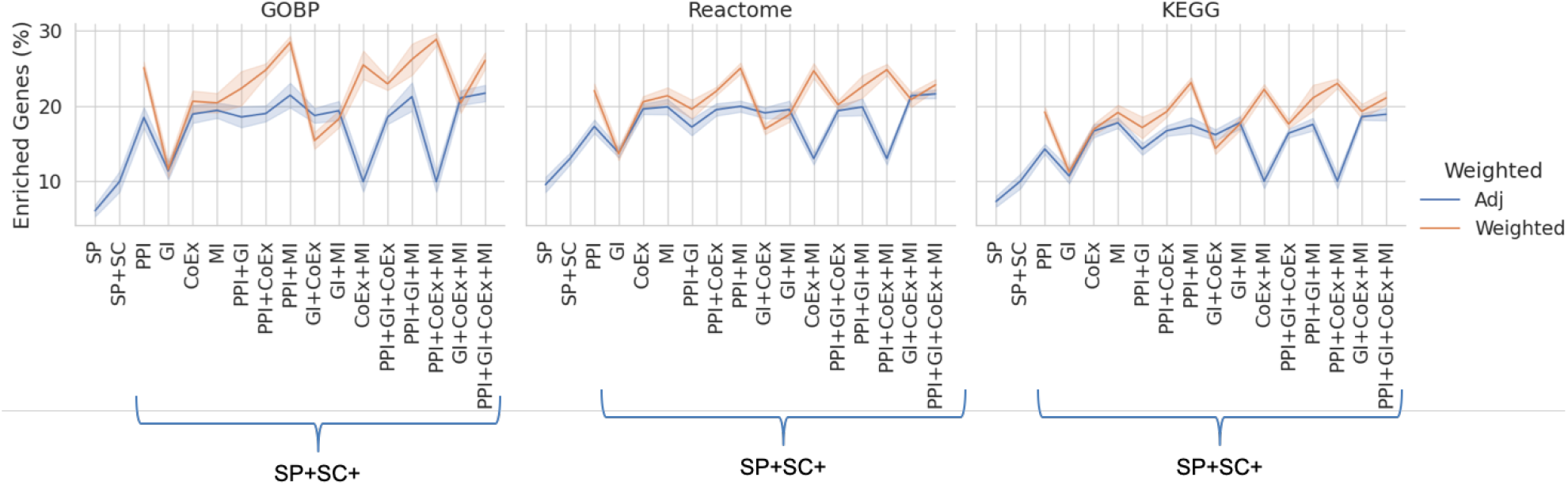
For different combinations of networks (x-axis) integrated with single-cell and spatial data with the iTissue data integration framework, percentage of enriched genes in GeneOntology Biological Process (left), Reactome Pathways (middle) and KEGG Pathways (right) using two representation for the molecular networks: adjacency matrix (blue) and weighted version (orange).

**Figure 2.**
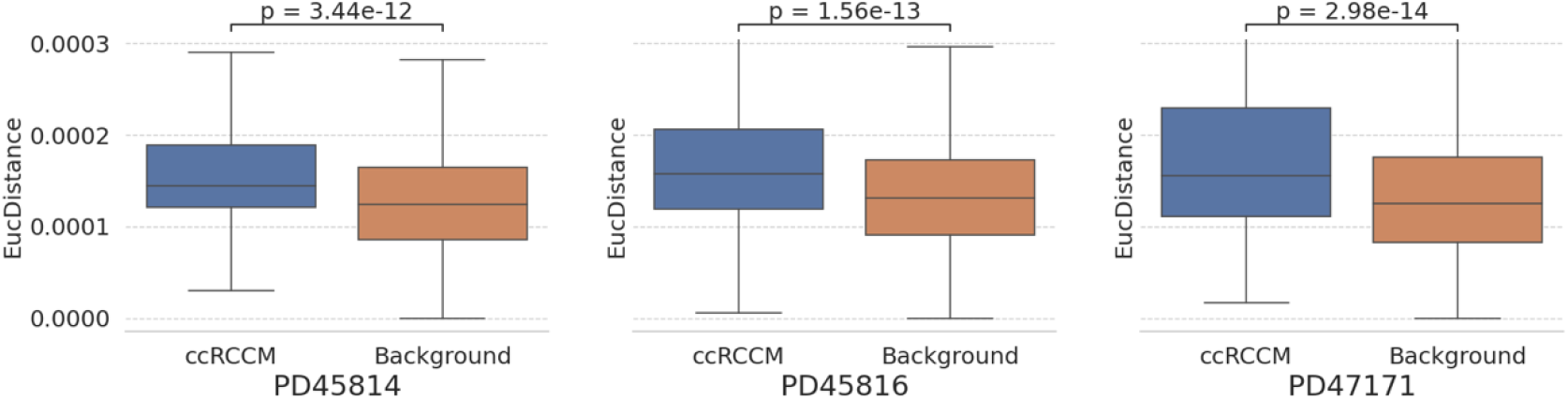
For the three patients, genes known to be related to clear cell renal cell carcinoma (ccRCC), in blue, have statistically significantly higher euclidean distance between the spaces created by the tumour-core and tumour-normal interface samples than the rest of the genes (Background), in orange. p-values obtained using Mann Whiteney U-test.

### Identifying relevant genes for metastatic cancer

To identify genes and mechanisms that differ in the different regions of a tumour, we compare the tumour-core and tumour-normal interface of 3 clear cell renal cell carcinoma (ccRCC) patients^10^ (see section “Spatial and single cell transcriptomic”). For each patient and sample (i.e., tumour-core and tumour-normal interface), we apply our iTissue data integration framework (see section “iTissue data Integration framework”) to integrate the spatial and single-cell data with the PPI and MI networks. We did not integrate CoEx and GI since we show that adding them in the integration do not result in a better functional organisation of the space (for more details, see section “Ablation Study”).

To assess if we can identify genes relevant for ccRCC using their movement between the spaces created by the iTissue, for each patient and sample (i.e., the tumour-core and tumour-normal interface), we compute the euclidean distance of each gene between the two spaces. Since the space created for each sample will have a different set of coordinates, to be able to compute the euclidean distance between genes of different spaces, we first compute the relative distance between the genes of the same space: *RelativeGeneEmbedding*(*RGE*) = *G* * *G*^*T*^. After computing the euclidean distance between each gene’s RGE of the spaces created by the tumour-core and tumour-normal interface samples, we compare the euclidean distances of the genes known to be related to ccRCC in DisGeNET database^11^ using Mann Whiteney U-test. We find that genes known to be related ccRCC have statistically significantly higher euclidean distance between the spaces created by the tumour-core and tumour-normal interface samples. Thus, other genes with high euclidean distance might also be relevant for ccRCC.

After showing that relevant genes for ccRCC have high euclidean distance between the spaces created by the tumour-core and tumour-normal interface samples, for each of the patients, we selected the top 100 with the highest euclidean distance. These three sets overlap in 50 genes (see Figure 3). Using the curated version of DisGeNET database, we validate that 26 of these 50 genes are known to be related to cancer. To investigate which specific mechanisms these 50 genes are participating in, we use enrichment analysis in GO-BP terms (for more details, see “Enrichment Analysis” section). Among all the enriched terms, we found that this set of genes are enriched in terms related to metastasis, such as, *positive regulation of endocytosis*, which is used by metastatic-suppressor genes to suppress metastasis^12^ and *regulation of fibroblast proliferation*, which play a key role in cancer invasion and metastasis^13^. Moreover, other enriched mechanisms, such as *regulation of cellular senescence*, which is a potential therapeutic target to limit metastasis formation^14^ and *Antigen processing and presentation*, which has been used in cancer immunotherapy^15^, suggesting that the identified genes could be used as potential treatments. These results show that using the euclidean distance between the spaces created by the tumour-core and tumour-normal interface samples for the three patients, we uncover a unique set of 50 genes that are involved in the metastatic process and that could be used as targets for drug repurposing.

**Figure 3.**
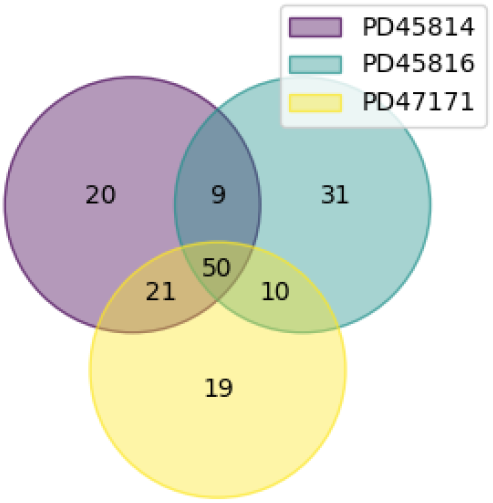
The top 100 with the highest movement of each patient overlap in 50 genes.

### Identified genes cluster the tumour-normal interface regions

To assess whether the identified genes are relevant for the three regions in the tumour-normal interface (healthy tissue, tumour tissue and interface), we cluster the spots in the Vissum array based on the distance of their embeddings to the embeddings of the identified genes in the gene space created by the iTissue. We cluster the spots using hierarchical clustering on the cosine distance between the spots embeddings and the genes embeddings (for more details, see “Clustering spots based on the identified genes” section). For two of the three patients (PD45816 and PD47171), the spots cluster in three big clusters (see Figure 4 and 5) and these correspond to the three regions of healthy tissue, tumour tissue and interface (see Figure 7 left and middle). For the patient PD45814, the majority of the spots are assigned to the same cluster (see Figure 6) and the three biggest clusters did not correspond to the three regions of healthy tissue, tumour tissue and interface (see Figure 7 right). These results show that, for two of the patients, the identified can separate the tumour-normal interface tissue into the three regions of healthy tissue, tumour tissue and interface.

**Figure 4.**
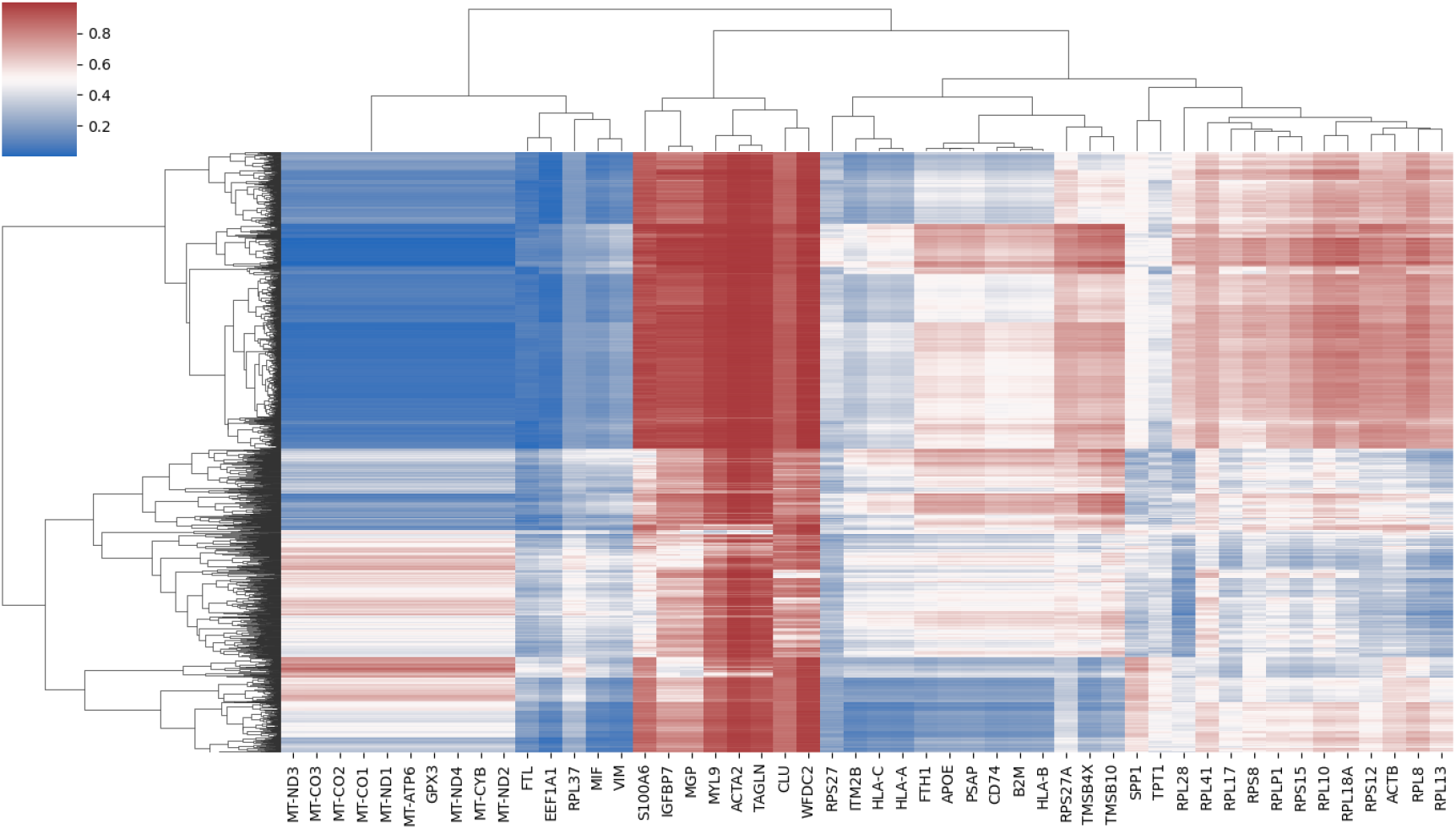
Hierarchical clustering of the spots using the cosine distance between the spots embeddings and the genes embeddings of the 50 genes with the highest movement for patient PD45816.

**Figure 5.**
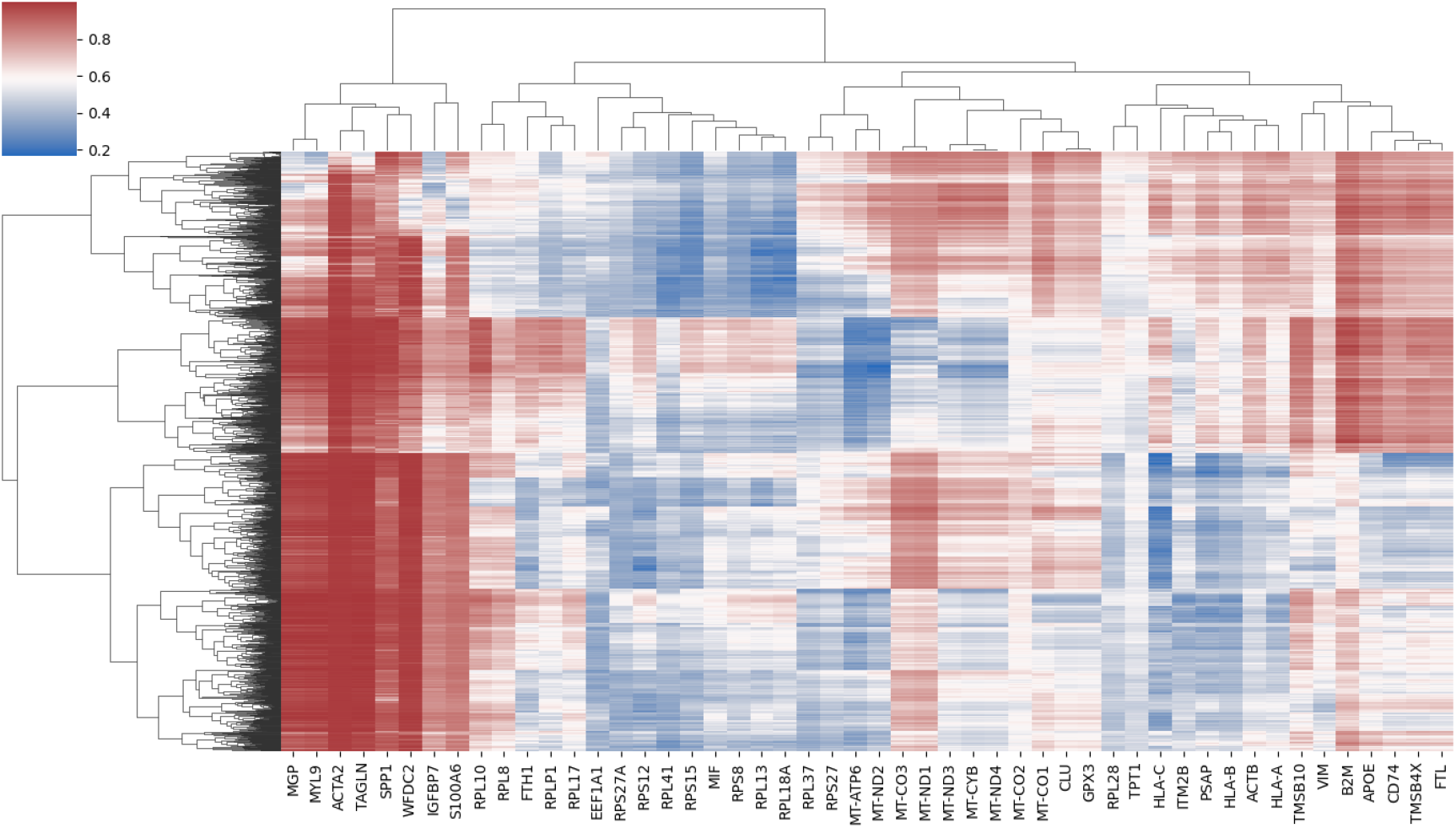
Hierarchical clustering of the spots using the cosine distance between the spots embeddings and the genes embeddings of the 50 genes with the highest movement for patient PD47171.

**Figure 6.**
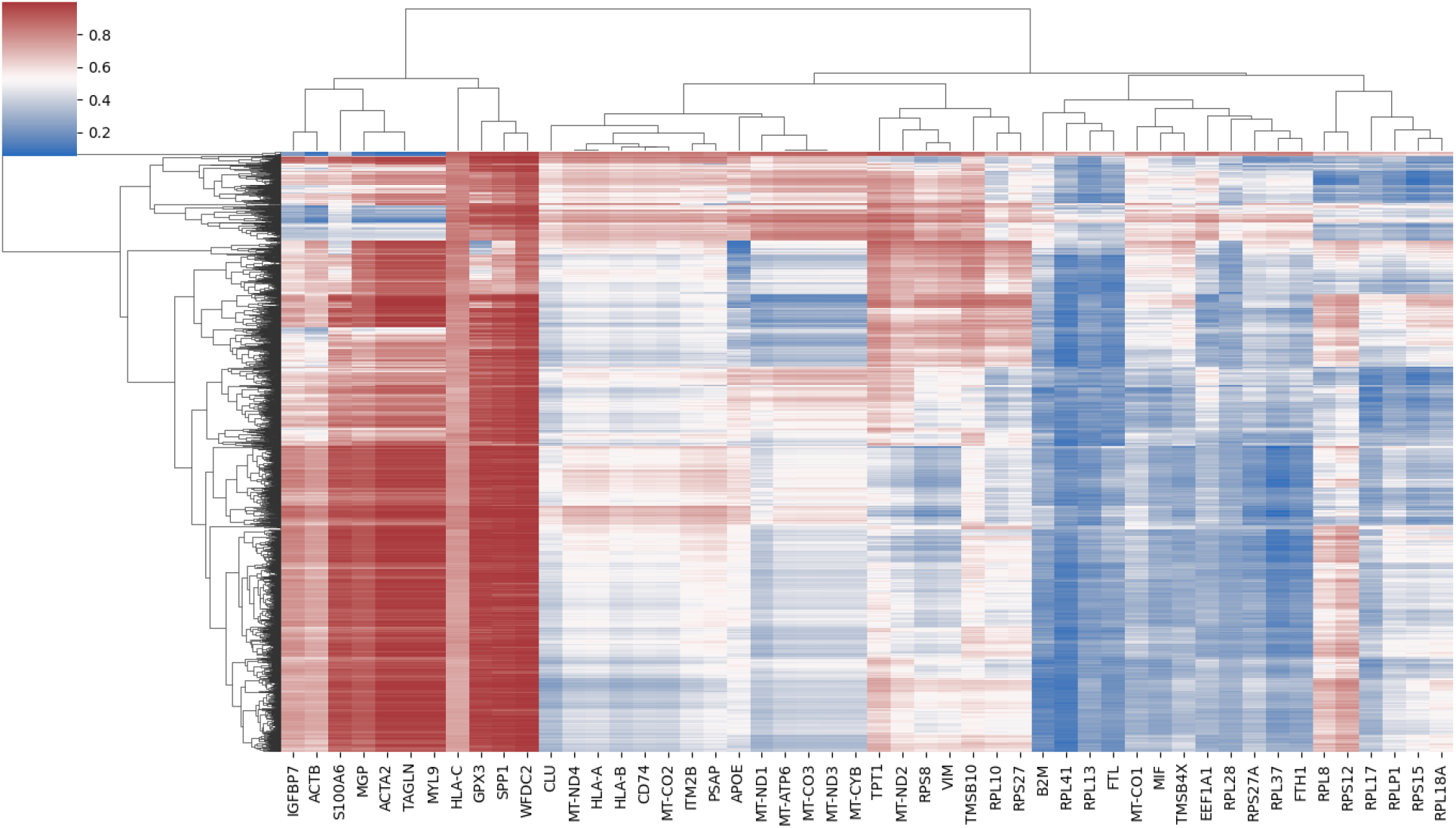
Hierarchical clustering of the spots using the cosine distance between the spots embeddings and the genes embeddings of the 50 genes with the highest movement for patient PD45814.

**Figure 7.**
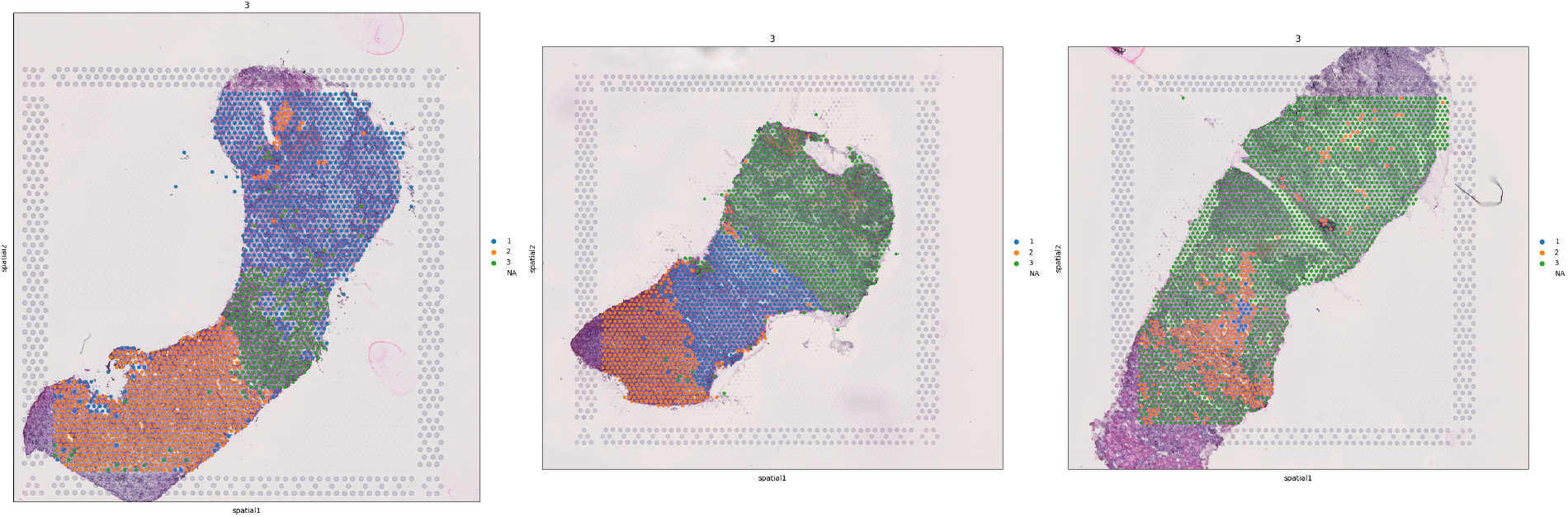
Clusters obtained using the hierarchical clustering define the three regions (healthy, tumour tissue and interface) for patients PD45816 (on the left) and PD47171 (in the middle). For patient PD45814 most of the spots correspond to the same clustering, and the clustering does not correspond to the regions.

**Figure 8.**
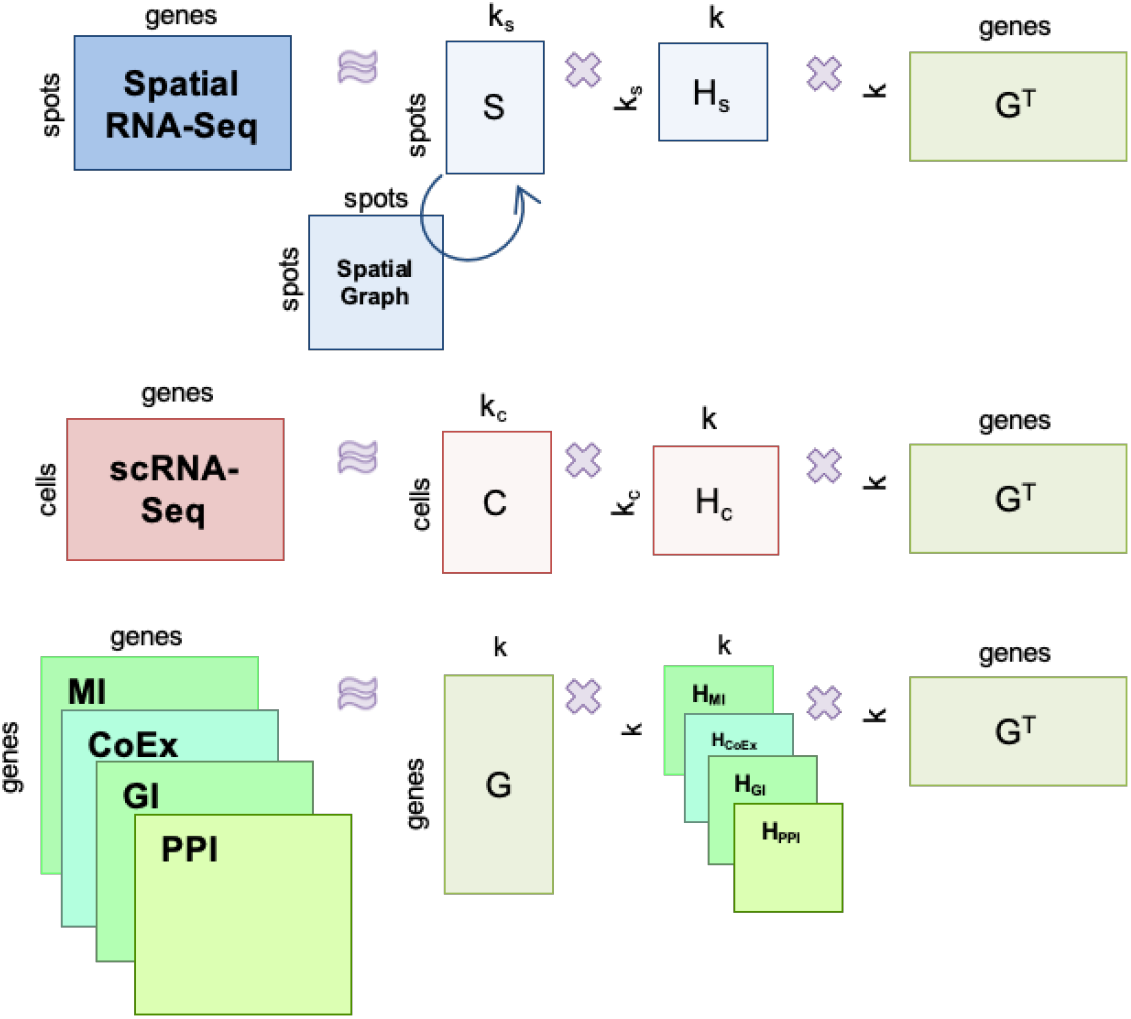
Illustration of the iTissue integration method. Graph-regularised non-negative matrix tri-factorisation (GNMTF) used for integrating the single-cell expression matrix, the spatial expression matrix, and all four matrices representing the molecular networks (PPI, GI, CoEx and MI). The matrix factor is shared across decompositions to simultaneously decompose all the matrices. The spatial information between the spots is incorporated into the data integration by using one regularisation term (illustrated by arcs with arrows).

## Discussion

In summary, we presented the iTissue framework to integrate spatial and single-cell transcriptomics data with the molecular interaction networks of PPI and MI. We apply iTissue to the tumour-core and tumour-normal interface samples of 3 clear cell renal cell carcinoma (ccRCC) patients. We showed that relevant genes for ccRCC have higher euclidean distance between their embedding in the spaces created by the tumour-core and tumour-normal interface samples. Thus, for each patient, we prioritise the top 100 genes with the highest euclidean distance between the gene embeddings obtained by these two samples. From these top 100 genes, we identify 50 genes that are shared between the 3 patients and validate that 27 out of the 50 are already known to be cancer related. Moreover, we validate that the identified genes are enriched in GO-BP relevant for metastatic mechanisms. Finally, we find that the identified genes can be used to cluster the tumour-normal interface in the main three regions (healthy tissue, tumour tissue and interface).

In this study, we uncover genes that are involved in metastatic mechanisms for ccRCC patients. However, the number of patients is limited to 3, so this result should be validated with more patients when data is available. Moreover, for two of the three patients, we cluster the tumour-normal interface in the three main regions (healthy tissue, tumour tissue and interface) using the identified genes. Further investigation is needed to assess why for the thrid patients the clusters obtained do not match the three main regions.

## Methods

### Data collection

#### Spatial and single cell transcriptomics

To identify relevant genes for the metastasis of the clear cell renal cell carcinoma (ccRCC), we use spatial and single-cell transcriptomics from 3 ccRCC patients^10^. From each patient, two samples were obtained, one from the core of the tumour and one from the tumour-normal interface. We create the single-cell expression matrix in which rows represent cells, columns represent genes, and an entry is the expression level. Similarly, we create the spatial expression matrix in which rows represent a spot in the Visium array, columns represent genes, and an entry is the expression level. To include the spatial information between the spots, we create the Spatial spot graph connecting each spot to all its neighbours.

#### Molecular networks

To integrate the single-cell and spatial data with prior knowledge, we collect four molecular networks from Homo sapiens. To create the protein-protein interaction (PPI) network, we collected all physical protein-protein interactions between human proteins from BioGRID v4.4.208^16^ that are experimentally validated using yeast two-hybrid, or affinity capture-based technologies; this resulted in 18,525 proteins (nodes) connected by 609,495 interactions (edges). To create the Gene interaction (GI) network, we collect all genetic interactions reported in BioGRID v4.4.208^16^, which resulted in 7,016 proteins (nodes) connected by 17,789 interactions (edges). To create the CoExpression (CoEx) network, we collect correlations between genes from CoexpressDB v.7.325^17^, the co-expression measure between two genes is standardised using zeta scores. To construct the COEX networks, in which nodes represent genes and edges represent the co-expressions between the genes, we selected the strongest co-expression values having a zeta score higher than or equal to 3, which is the usual practice, this resulted in resulted in 15,130 proteins (nodes) connected by 14,63280 interactions (edges). Finally, we construct the Metabolic Interaction (MI) network by connecting genes participating in the same metabolic pathways in KEGG^9^, resulting in 3,415 proteins (nodes) connected by 465,112 interactions (edges).

Following^18^, we represent the PPI, GI and MI networks with their Positive Pointwise Mutual Information (PPMI) matrices. These matrices quantify how frequently any two nodes in the corresponding PPI, GI and MI networks co-occur in a random walk compared to what would be expected if the occurrences of the nodes were independent. We use the DeepWalk closed formula with its default settings to compute the PPMI matrix (see Equation 1)

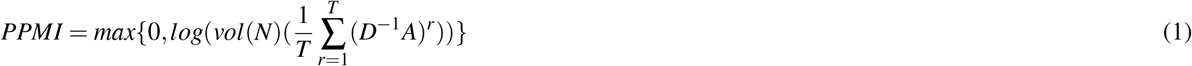

where A is the adjacency matrix of the network N, D is the diagonal matrix of A, vol(N) is the volume of N, *T* = 10 is the length of the random walk.

To have a corresponding weighted matrix for the CoEx network, we took as weights the normalised correlation scores.

### iTissue data integration framework

To integrate the spatial and single-cell transcriptomic with molecular interaction networks, we used Graph-regularized Non-negative Matrix Tri-Factorization (GNMTF), which approximates a high-dimensional data matrix as a product of three low-dimensional, non-negative matrices. We decompose the single-cell expression matrix in three matrices *S, H*_*s*_ and *G*, the spatial expression matrix in three matrices *C, H*_*c*_ and *G*, and all four matrices representing the molecular networks in three matrices *G, H*_*i*_ where *i* = {PPI, GI, COEx, MI} and *G*. The matrix factor *G* is shared across decompositions to simultaneously decompose the all the matrices. The spatial information between the spots is added to the decomposition through graph regularisation penalties. We use these graph regularisation penalties to favour embedding close in space the spots that are neighbours in the spatial spot graph (see Figure 8)

To extract knowledge from the decomposition, we embedded all the entities, spots, cells, and genes in the gene space. We obtain the spot embeddings by multiplying *S* × *H*_*s*_, and the cell embedding by multiplying *C* × *H*_*c*_. For the gene embeddings, we directly used the matrix factor *G*.

### Extracting clusters of genes

The matrix factor *G*^*g*×*k*^, from the GNMTF decomposition, is the cluster indicators of genes; based on their entries, *g* genes are assigned to *k* clusters. In particular, the hard clustering procedure of Brunet *et al*. 2004^19^, was used to cluster the genes of the matrix factor *G*^*g*×*k*^. The columns of *G*^*g*×*k*^ correspond to the *k* clusters, and each gene is assigned to the cluster that has the largest entry in the gene row.

### Enrichment Analysis

To assess the quality of the obtained clusters of genes, we computed the enrichment of biological annotations in the clusters. For each gene (or equivalently, protein, as a gene product) in the network, we use the most specific experimentally validated annotations of Biological Process (BP) present in the Gene Ontology (GO)^7^. The probability that an annotation is enriched in a cluster was computed by using a hypergeometric test, i.e., sampling without replacement strategy shown in equation 2:

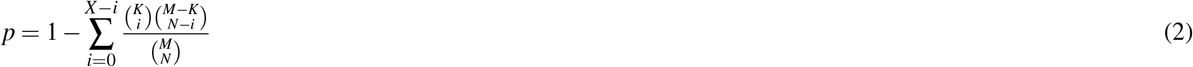

where *N* is the number of annotated genes in the cluster, *X* is the number of genes in the cluster that are annotated with the given annotation, *M* is the number of annotated genes in the network and *K* is the number of genes in the network that are annotated with the annotation in question. Annotations with a Benjamini-Hochberg adjusted p-value^20^ of *p* ≤ 0.05 were considered to be statistically significant enriched.

### Clustering spots based on identified genes

To assess whether the identified genes are relevant for the three regions in the tumour-normal interface (healthy tissue, tumour tissue and interface), we cluster the spots in the Vissum array based on the distance of their embeddings to the embeddings of the identified genes in the gene space created by the iTissue. First, we obtain the spots embeddings in the gene space by multiplying matrix factors obtained by the decomposition *SpotEmbeddings*(*SE*) = *S* × *H*_*s*_. Then, we compute the cosine distance between the spots embeddings and the genes embeddings and cluster the spots according to it using Hierarchical clustering.

## Acknowledgements

This project has received funding from Mohamed bin Zayed University of Artificial Intelligence, the European Research Council (ERC) Consolidator Grant 770827, the Spanish State Research Agency and the Ministry of Science and Innovation MCIN grant PID2022-141920NB-I00/AEI/10.13039/ 501100011033 FEDER, UE, and the Department of Research and Universities of the Generalitat de Catalunya code 2021 SGR 01536.

## References

1. Mihajlovic K, Ceddia G, Malod-Dognin N, Novak G, Kyriakis D, Skupin A, et al. Multi-omics integration of scRNA-seq time series data predicts new intervention points for Parkinson’s disease. bioRxiv. preprint. 2023;12:2023.

2. Mihajlović K, Malod-Dognin N, Ameli C, Skupin A, Pržulj N. MONFIT: multi-omics factorization-based integration of time-series data sheds light on Parkinson’s disease. NAR Molecular Medicine. 2024;1(4):ugae012.

3. Yang J, Yang S, Fu Y, Li X, Huang T. Non-negative graph embedding. In: 2008 IEEE Conference on Computer Vision and Pattern Recognition. IEEE; 2008. p. 1–8.

4. Huizing GJ, Deutschmann IM, Peyré G, Cantini L. Paired single-cell multi-omics data integration with Mowgli. Nature Communications. 2023;14(1):7711.

5. Argelaguet R, Arnol D, Bredikhin D, Deloro Y, Velten B, Marioni JC, et al. MOFA+: a statistical framework for comprehensive integration of multi-modal single-cell data. Genome biology. 2020;21:1–17.

6. Cai D, He X, Han J, Huang TS. Graph regularized nonnegative matrix factorization for data representation. EEE Transactions on Pattern Analysis and Machine Intelligence,. 2010;33(8):1548–60.

7. Ashburner M, Ball CA, Blake JA, Botstein D, Butler H, Cherry JM, et al. Gene ontology: Tool for the unification of biology. Nature Genetics. 2000 may;25(1):25–9.

8. Jassal B, Matthews L, Viteri G, Gong C, Lorente P, Fabregat A, et al. The reactome pathway knowledgebase. Nucleic Acids Research. 2019 11;48(D1):D498-503.

9. Kanehisa M, Furumichi M, Tanabe M, Sato Y, Morishima K. KEGG: New perspectives on genomes, pathways, diseases and drugs. Nucleic Acids Research. 2017 jan;45(D1):D353-61.

10. Li R, Ferdinand JR, Loudon KW, Bowyer GS, Laidlaw S, Muyas F, et al. Mapping single-cell transcriptomes in the intra-tumoral and associated territories of kidney cancer. Cancer cell. 2022;40(12):1583–99.

11. Piñero J, Ramírez-Anguita JM, Saüch-Pitarch J, Ronzano F, Centeno E, Sanz F, et al. The DisGeNET knowledge platform for disease genomics: 2019 update. Nucleic Acids Research. 2019 11;48(D1):D845-55.

12. Khan I, Steeg PS. Endocytosis: A pivotal pathway for regulating metastasis. British journal of cancer. 2021;124(1):66–75.

13. Asif PJ, Longobardi C, Hahne M, Medema JP. The role of cancer-associated fibroblasts in cancer invasion and metastasis. Cancers. 2021;13(18):4720.

14. Faggioli F, Velarde MC, Wiley CD. Cellular senescence, a novel area of investigation for metastatic diseases. Cells. 2023;12(6):860.

15. Lee MY, Jeon JW, Sievers C, Allen CT. Antigen processing and presentation in cancer immunotherapy. Journal for immunotherapy of cancer. 2020;8(2):e001111.

16. Oughtred R, Stark C, Breitkreutz BJ, Rust J, Boucher L, Chang C, et al. The BioGRID interaction database: 2019 update. Nucleic Acids Research. 2019 jan;47(D1):D529-41.

17. Obayashi T, Kagaya Y, Aoki Y, Tadaka S, Kinoshita K. COXPRESdb v7: a gene coexpression database for 11 animal species supported by 23 coexpression platforms for technical evaluation and evolutionary inference. Nucleic acids research. 2019;47(D1):D55-62.

18. Doria-Belenguer S, Xenos A, Ceddia G, Malod-Dognin N, Pržulj N. The axes of biology: a novel axes-based network embedding paradigm to decipher the functional mechanisms of the cell. Bioinformatics Advances. 2024;4(1):vbae075.

19. Brunet JP, Tamayo P, Golub TR, Mesirov JP. Metagenes and molecular pattern discovery using matrix factorization. Proceedings of the National Academy of Sciences of the United States of America. 2004 mar;101(12):4164–9.

20. Benjamini Y, Hochberg Y. Controlling the False Discovery Rate: A Practical and Powerful Approach to Multiple Testing. Journal of the Royal Statistical Society: Series B (Methodological). 1995 jan;57(1):289–300.

